# The parafascicular nucleus of the thalamus orchestrates coordinated skeletomotor, autonomic, and aversive state transitions

**DOI:** 10.64898/2026.02.19.706912

**Authors:** Marina Roshchina, Henry H. Yin

**Affiliations:** Department of Psychology and Neuroscience; Duke University; Durham, NC, 27708; USA.; Department of Neurobiology; Duke University School of Medicine; Durham, NC, 27708; USA

**Keywords:** parafascicular nucleus of the thalamus, intralaminar thalamus, facial expression, autonomic function, valence, optogenetics

## Abstract

The parafascicular nucleus (Pf), part of the intralaminar thalamic nuclei, has been implicated in diverse functions such as attention, nociception, and behavioral flexibility, yet its precise contributions to behavior remain poorly defined. In this study, we used optogenetics in male and female mice to study the role of Pf projection neurons using high-resolution and continuous measures to quantify both skeletomotor and autonomic behavioral outputs as well as motivational valence. We showed that selective Pf stimulation resulted in wide-ranging effects, including ipsiversive turning, facial and whisker movements, pupil constriction, and heart rate reduction. Finally, we found that Pf stimulation could be highly aversive, as mice showed strong place aversion to areas where stimulation was delivered. Together our results indicate that Pf outputs can ultimately influence both skeletomotor outputs like turning and autonomic outputs like pupil constriction. The latter parasympathetic responses may be directly related to the role of the Pf in regulating motivational and emotional valence. These findings broaden our understanding of Pf function and suggest it may serve as a major hub for the integration of behavioral state feedback and top-down command of a variety of effectors.

**Significance statement:** This study demonstrates that the parafascicular nucleus (Pf) of the thalamus plays a critical role in coordinating both skeletomotor and autonomic responses, as evidenced by optogenetic stimulation in mice triggering rapid ipsilateral turning, facial movements, pupil constriction, and heart rate reduction. The findings reveal that Pf activation induces an aversive state, highlighting its involvement in motivational valence. These insights expand our understanding of Pf’s top-down regulatory influence on behavior.

---

The parafascicular nucleus (Pf) of the thalamus is a part of the intralaminar thalamic nuclei, which are traditionally considered a relay station between brainstem and the cortico-basal ganglia circuits. Studies have shown that the Pf is not a homogenous nucleus but comprises different neuronal subpopulations with a topographical organization (Feger et al., 1994; Mandelbaum et al., 2019). The Pf receives projections from many brain regions including the superior colliculus, trigeminal nuclei, vestibular nuclei, nucleus coeruleus, dorsal nucleus of Raphe, and parabrachial nucleus (Comans and Snow, 1981; Cornwall and Phillipson, 1988; Chen et al., 1992). As a main source of thalamostriatal projections, it has a strong influence on basal ganglia activity, and previous work mostly focused on its projections to the dorsal striatum (Gonzalo-Martín et al., 2024). But it also projects to many other areas such as the subthalamic nucleus and cerebral cortex (Groenewegen and Berendse, 1994; Kumar et al., 2023).

Perhaps reflecting this anatomical complexity, Pf has been implicated in different functions such as nociception (Peschanski et al., 1981; Weigel et al., 2004), action selection (Stayte et al., 2021), action initiation and steering behavior (Watson et al., 2021; Fallon et al., 2023), and behavioral flexibility (Bradfield et al., 2013). Recent work has shown that Pf neurons can encode movement kinematics during turning, and stimulation of Pf can generate ipsiversive turning behavior (Fallon et al., 2023). These findings that Pf projections can directly influence skeletomotor output, but the extent of this contribution remains unclear. For example, it is unknown whether Pf output mainly influences behavior related to turning and orienting, or whether it can also target other types of behavioral outputs.

In the present study, we performed a detailed analysis of motor and autonomic responses to optogenetic stimulation of the main projection neurons (Vglut2^+^) in the Pf. In addition to replicating previously observed turning behavior, we report a number of new findings on the contribution of the Pf in behavior. We found that Pf stimulation resulted in facial motor responses, including movements of the nose and whiskers. Additionally, we found that Pf stimulation had a strong and persistent effect on autonomic outputs. It could produce parasympathetic responses such as pupil constriction and a decrease in heart rate. These responses are associated with strong aversion, as mice quickly learn to avoid the area associated with Pf stimulation.

## Materials and Methods

### Mice

All experimental and surgical procedures were performed in accordance with standard protocols and approved by the Duke University Institutional Animal Care and Use Committee. Mice were at least 10 weeks old at the start of the experiment. All mice were housed in groups of 2-5 per cage with free access to food and water and maintained on a 12/12 h light/dark cycle. Experiments were performed during the light phase. Data were collected from 21 mice (15 males and 6 females. Vglut2-ires-Cre (Slc17a6tm2(cre)Lowl) and C57BL/6J lines. Vglut2-ires-Cre mice were randomly assigned to experimental (opsin, n=12) and control (n=6) groups. In addition, 3 C57BL/6J mice were also used as controls (total number of Control animals = 9).

### Viral Vectors

To manipulate neural activity optogenetically, we used adeno-associated virus (AAVs) to express CoChR or a control fluorophore YFP. For the experimental group, we injected AAV9-EF1a-DIO-CoChR-Kv2.1-GP2A-eGFP (Duke University Vector Core (0.9×10^12 GC/ml)). This construct allows Cre-dependent expression of the excitatory opsin ChR2 fused to enhanced yellow fluorescent protein (EYFP). This vector contains the excitatory opsin soma-targeted Chloromonas oogama Channelrhodopsin that can be activated by blue light (Forli et al., 2021). For the control mice pAAV-Ef1a-DIO EYFP (Addgene #27056; 1.6*10^13 GC/ml). In case of C57BL/6J control animals we injected AAV9-EF1a-DIO-CoChR-Kv2.1-GP2A-eGFP virus.

### Surgery

Mice were anesthetized with 3% and maintained at 1.5% isoflurane in oxygen at a rate of 1 l/min during surgical procedures. They were fixed in a stereotactic frame (David Kopf Instruments, Tujunga, CA) and administered Meloxicam (2 mg/kg) and bupivacaine (0.20 mL) before surgical incisions. Separate craniotomies for virus and fibers were made bilaterally. Virus (60 nl) was injected at 1nl/s with a microinjector (Nanoject 3000, Drummond Scientific) through a glass pipette into Pf (relative to Bregma: AP: -2.3 mm; ML±0.65-0.75 mm; DV: -3.6/-3.2 mm). After 5 minutes post-injection period custom-made optic fibers (0.22 NA, 105-µm core, ∼75-85% transmittance) were lowered into Pf (relative to Bregma: AP: -2.3 mm; ML±1.5 mm; DV: - 2.75 mm, at 15°) and fixed with acrylic cement (Stoelting) with skull screws and the metal custom-made head bars for head fixation. Mice were allowed to recover for three weeks before the start of experiments.

### Optogenetic stimulation in Open Field

For open field stimulation experiments mice were unilaterally connected to a 473-nm diode-pumped solid-state laser (Shanghai Laser) via custom-made fiber-optic cables and rotatory joint (Doric) and allowed to explore a circular arena (diameter – 44 cm) for 10-15 minutes. Behavior was recorded from the top with a BlackFly video camera (TeledyneVision Solutions) at 50 fps. Video recording, control of the laser pulses delivery and collection of timestamps were managed through the Bonsai software (Lopes et al., 2015) and Arduino board with custom script. To excite Vglut2+ neurons, trains of 10 ms pulses with a frequency of 20 Hz were delivered. The laser power at the tip of the optical fiber was measured (PM120VA, Thorlabs) before each session and was set to 2, 4 or 6 mW. Stimulation was manually controlled through the Bonsai software, with stimuli presented in a random order, each lasting 2 seconds with at least 10 seconds between presentations. Deeplabcut (DLC, version 2.3.9) was used for offline detection and tracking of head and body centers (Mathis et al., 2018). Custom python script was used for calculation of instant speed for each DLC marker and angular velocity of the body to head vector.

### Facial expression measure

Recording of facial expression was performed in mice head-fixed in custom restraining setup by metal bars attached to the cement cap. Mouse face profile was recorded under dark conditions (less than 1 lux) under infrared light with BlackFly video camera (TeledyneVision Solutions) at 100 fps. Laser was controlled manually through an Arduino Uno serial controller. Cerebus Neural Signal Processor (Blackrock Microsystems) was used to record frame signal from camera and control signal stamps for laser from Arduino. Stimulation of Pf neurons was performed using trains of laser pulses (10 ms pulse width, 1 second train length). The stimulation protocol involved three different experimental conditions: 1) pulse frequency was maintained at 20 Hz while the laser power was varied (2, 4 or 6 mW); 2) power was fixed at 6 mW, and the pulse frequency was adjusted (2, 5, 10 or 20 Hz); 3) power and frequency were fixed at 6mW, 20Hz and the duration of the stimulation was varied (250ms = 5 pulses, 500ms = 10 pulses, 750ms = 15 pulses, 1000 ms = 20 pulses).

To describe the stimulation-evoked changes in facial expression, the whisker pad movement was calculated based on changes in position of the base of one macrovibrissa. Whisker velocity data was not standardized because of high similarity between mice. The nose position was determined by the angle between horizontal line and a nose vector (defined between nose base and tip). One control mouse developed glaucoma on one eye after the start of the experiment, in this animal data from only one pupil was recorded.

### Heart Rate measure

To measure heart rate 6 CoChR and 4 control mice were implanted with custom made Electrocardiogram (ECG) electrodes. Two electrodes were used for the ECG signal, while the third electrode was used as the reference. During surgery two skin incisions were made on the upper right and lower left parts of the rib cage. The ECG electrode connector was secured on the cement cape together with optic fibers and two wires were passed subcutaneously from head to the incisions on torso and sutured onto muscle tissue. The reference electrode was passed subcutaneously to the middle of the back and left free.

ECG optogenetic experiment was performed on unrestrained mice placed in small plastic beaker. For optogenetic stimulation the same setup as described in Facial recording experiment was used. For ECG recording the head connector was attached to the SparkFun Heart Rate Monitor (AD8232, SparkFun Electronics) which transmitted the differentiated ECG signal to the Cerebus Neural Signal Processor (Blackrock Microsystems). Raw ECG signal was collected at 2 kHz together with the video frame timestamps and laser control timestamps. Neuroexplorer was used for exporting the raw ECG data and laser timestamps. Raw signal detection and extraction of heartbeats from the raw signal were performed with a custom Python script. A highpass filter was applied to remove slow baseline fluctuation. Heart beats were detected as local maximums with ‘find peaks’ python function.

### Real-time place avoidance (RTPA)

Real-time place avoidance test was performed in rectangle plastic arena (35×28×23 cm). The video recording and stimulation setup was similar to the open field experiment. The Bonsai software was used for control on video recording, position tracking, laser delivery and timestamps collecting. An RTPA test consists of the three epochs: 5-min pretest (Before), 5-min laser stimulation (Laser) and 5-min after laser (After). During the Before epoch mice were allowed to freely explore the entire arena and the preferred half of the arena was determined. For the next epoch the preferred half was assigned as ‘Laser’, and a rectangular ROI was selected around that part of the arena. During the Laser epoch the laser was on every time mouse entered the defined ROI (10ms pulses, delivered at 20 Hz with 6 mW power). The laser stayed on while the mouse was in the ROI. After 5 minutes, the laser was turned off and the next epoch (‘After’) started. Videos recorded during RTPA were analyzed with DeepLabCut and custom python script in the same manner as in open field experiment.

### Histology

Mice were transcardially perfused with 0.1 M phosphate buffered saline and then 4% paraformaldehyde. Brains were extracted and submerged in 4% paraformaldehyde 1-2 days and then in 30% sucrose for 2-3 days. After sinking, brains were sliced coronally at 60 μm with the cryostat (Leica CM1850). For position verification sections were immunostained against GFP and NeuN. On day 1 sections were rinsed in 0.1 M PBS for 30 min before being placed in a blocking solution containing 5% goat and donkey serum and 0.25% Triton X-100 at room temperature for 1 hour. Then all sections were incubated with primary antibodies [polyclonal chicken anti–green fluorescent protein (GFP), 1:500 dilution; Abcam, catalog no. ab13970 and monoclonal rabbit anti-NeuN, 1:500 dilution; Abcam, catalog no. ab177487] overnight at 4°C. On day 2 sections were rinsed in PBS for 30 min before being placed in a secondary for enhancing YFP signal (goat anti-chicken Alexa Fluor 488, 1:500 dilution; Thermo Fisher Scientific, catalog no. A-11039) and labeling neuronal nuclei (donkey anti-rabbit Alexa Fluor 594, 1:500 dilution; Jackson ImmunoReserach, catalog no. 711-585-152) for 2 hours at room temperature. After staining sections were mounted with an aqueous fluoroshield DAPI mounting medium (Abcam, Ab104139) and imaged (ZEISS 220 Axio Zoom.V16, SCR_016980). Data from 3 stimulation sides were excluded from the analysis because the optic fiber tips were outside of the Pf.

### Statistical Analysis

All statistical tests (t-test, ANOVA, mixed effects analysis) were performed in GraphPad Prism (GraphPad Software).

## Results

### Optogenetic Excitation of Vglut2^+^-Pf Neurons Initiates Fast Ipsilateral Turning

To examine how the parafascicular nucleus (Pf) contributes to movement control during locomotion, we injected Cre-dependent CoChR into the Pf of Vglut2-ires-Cre mice and bilaterally implanted optic fibers (Figure 1A-C). All experiments were based on the unilateral stimulation, with the effects of each stimulation site (hemisphere) analyzed independently.

**Figure 1.**
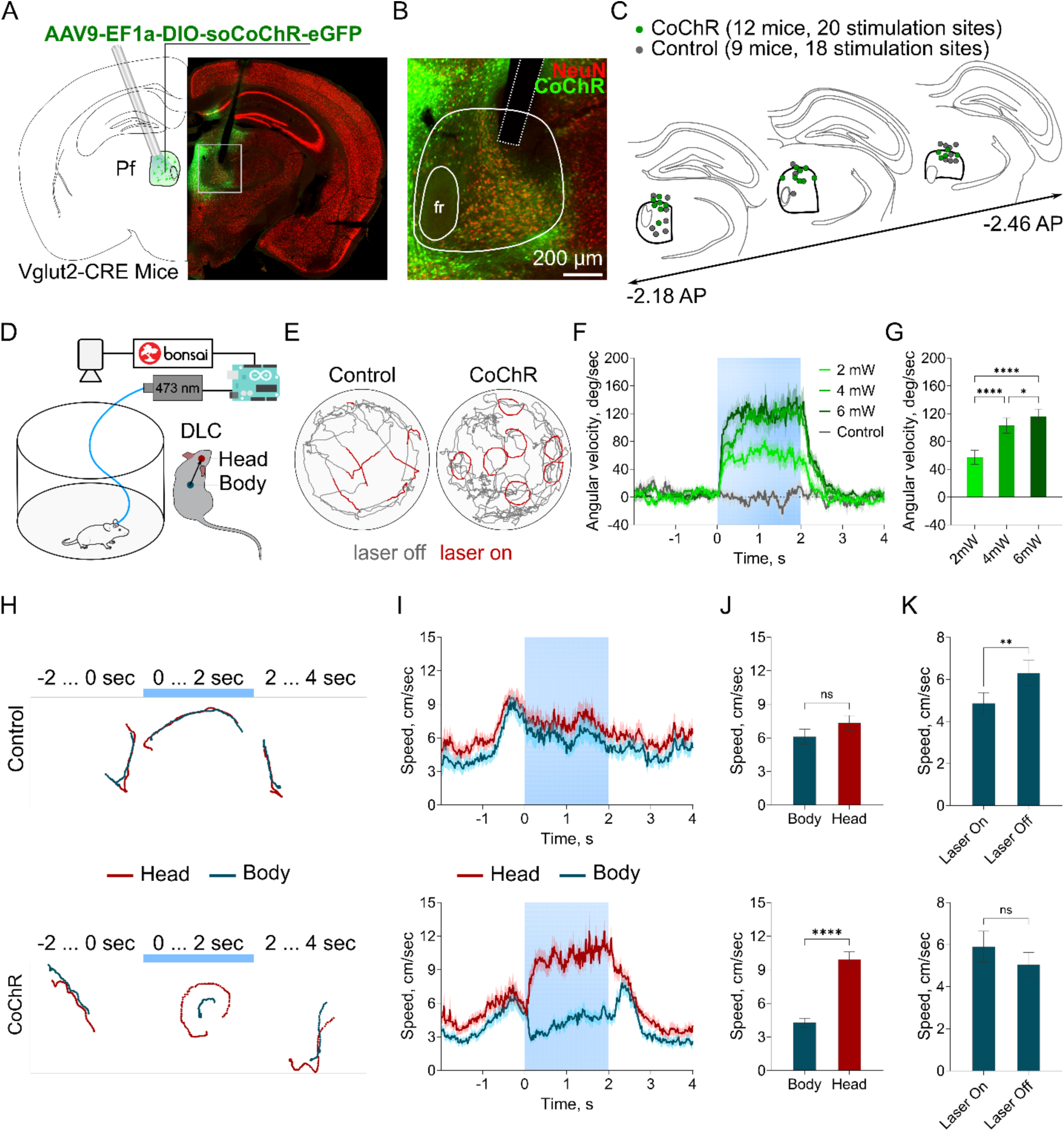
Unilateral excitation of Vlgut2^+^ Pf neurons initiates ipsilateral turns. A) Schematic of the surgery strategy and representative histology for the Cre-dependent CoChR expression and placement of optic fibers in Pf. B) Magnified view (white rectangle in A) showing the CoChR expression level and fiber tip relative to Pf. C) Optic fiber tip locations for CoChR (n=12, Vglut2^+^-CoChR-Pf) and control (n=9, 6 Vglut2^+^-eYFP-Pf and 3 C57Bl6-CoChR-Pf) animals. D) Schema of the Open Field experiment. E) Representative head trajectories during Open Field experiment. Left – control animal, right – CoChR expressing animal. Red – path during laser stimulation. F) Instant angular velocity increases during laser presentation in CoChR mice. Different shades of green shows different laser power in CoChR mice, grey – control mice at 6mW laser power. Blue bar indicates optogenetic stimulation. G) Average angular velocity during 2s laser presentation increases with laser power increase. H) Representative trajectory of head (red) and body (blue) 2 seconds before, 2 seconds of laser presentation (marked with blue line) and 2 seconds after of the single trial. I) Instant speed of the head and body did not differ in control animals but show noticeable difference in CoChR mice. J) Average speed of the head is higher than the body speed in CoChR mice but not in Control. Average speed calculated over 2 second of laser presentation. K) Average body speed increases after the laser end in CoChR mice, but not in Control mice. Average speed of the body calculating during the last 1 second of laser presentation (laser on) and 1 second after laser end (laser off). (for H, I, J, K – up – control mice, down – CoChR mice). All data represented as mean ± SEM, * - p< 0.0332, ** - p < 0.0021, **** - p < 0.0001, ns – p > 0.05.

We first stimulated Vglut2^+^-Pf neurons at 20 Hz for 2 seconds during open-field exploration (Figure 1D). Using DeepLabCut for offline tracking, we measured the movements of the head and body centers, calculating the speed of each part and the angular velocity of the vector between the head and body. Consistent with previous reports, unilateral optogenetic stimulation of Vglut2^+^-Pf reliably induced rapid ipsilateral turns which are reflected as rapid increase in angular velocity (Figure 1E-F). The increase in angular velocity started soon after the laser onset and remained until laser offset, for the 6mW stimulation trials, the latency from laser onset to the start of rotation was 98 ± 56 ms (Figure 5K). Control animals showed no behavioral change (cumulative angle during laser: 0.4 ± 24.4 degrees). Increasing laser power from 2 to 6 mW resulted in progressively larger angular velocities (Figure 1F-G). A repeated-measures ANOVA revealed a significant main effect of laser power on average angular velocity (RM ANOVA, F_(1.545, 29.35)_ = 39.12, p < 0.0001; post hoc Tukey 2mW vs 4mW p < 0.0001, 2mW vs 6mW p < 0.0001, 4mW vs 6mW p=0.04), with higher stimulation power producing faster turns.

Detailed analysis of head and body velocities revealed a consistent kinematic pattern. Following laser onset, in CoChR mice head velocity increased while body velocity either decreased or remained low during the stimulation period (Figure 1H-I). As a result, average head velocity during stimulation was significantly higher than average body velocity in CoChR mice (head - 9.9 ± 3.0 cm/s, body - 4.3 ± 1.8 cm/s; t-test, p < 0.0001, Figure 1J). This pattern indicates that observed increase in angular velocity is primary driven by rapid head reorientation, which often occurs independently of, and sometimes interrupts, ongoing locomotion. In contrast, control animals showed no such dissociation (Figure 1H-I). Head and body velocities remained closely matched during the stimulation period (body: 6.1 ± 2.3 cm/s, head: 7.3±2.4 cm/s; paired t-test, p = 0.2, Figure 1J). In addition to the effects during stimulation, we observed a brief increase in body velocity immediately after the laser offset in CoChR, but not in control mice (Figure 1K). This post-stimulation burst lasted approximately 0.5 s and was indicated by a significant increase in body speed (laser on: 4.9 ± 2.2 cm/s; laser off: 6.3 ± 2.7 cm/s; paired t-test, p = 0.004).

### Stimulation of Vglut2^+^-Pf Neurons is aversive in a Real-Time Place Avoidance Task

While open-field exploration experiments revealed mostly motor effects, we also observed behavioral patterns indicating the negative valence such as activity reduction, increased grooming and thigmotaxis. Given the recent evidence of the Pf association with the avoidance-related circuits(Chen et al., 2025), we next directly tested the valence of the optogenetic activation of Vglut2^+^-Pf neurons using a real-time place avoidance (RTPA) task (Figure 2A, C). Mice first explored a rectangular arena for 5 minutes, and using the Bonsai online tracking module, we identified their preferred half based on time spent. We then designated the preferred half as the “Laser” half and the other as the “Safe” half. For the next 5 minutes, mice received constant laser stimulation (6 mW, 20 Hz, 10ms pulse width) whenever they entered the “Laser” half, followed by 5 minutes of exploration without stimulation. We assessed avoidance by calculating the percent of time spent in each half and found that CoChR mice spent significantly less time in the “Laser” half during and after laser presentation than before or than control mice (Figure 2G). Two way ANOVA shown a significant interaction between Epoch (Before, Laser and After) and Group (CoChR and Control) (F_(1.513, 45.38)_ = 24.52, p<0.001), indicating that avoidance of the stimulation-related half was specific for CoChR mice. Post hoc comparison confirmed that CoChR mice spent significantly less time in “Laser” half during Laser epoch and After epoch than in Before epoch (CoChR Before vs. Laser p < 0.001, CoChR Before vs. After p < 0.001). During the Laser epoch CoChR mice not only avoided the “Laser” half but also moved less in the “Safe” half (average speed 2.5 ± 0.8 cm/s) compared to control mice (4.1 ± 1.4 cm/s; two-way repeated measures ANOVA, Group effect F_(1,30)_ = 9.90, p = 0.0037; post hoc Tukey CoChR vs. Control, ‘Before’ p = 0.18, ‘Laser’ p = 0.002, ‘After’ p = 0.07, Figure 2H). Total traveled distance was also decreased in CoChR mice during the ‘Laser’ epoch compared to control mice (CoChR - 717.7 ± 211.7 cm; Control - 998.0 ± 196.8 cm; unpaired t-test, p = 0.0008, Figure 2I), consistent with a negative-valence state and/or defensive suppression of locomotion. Importantly, CoChR mice still entered the Laser half but exited rapidly after the stimulation onset, consistent with an aversive contingency rather than inability to enter.

**Figure 2.**
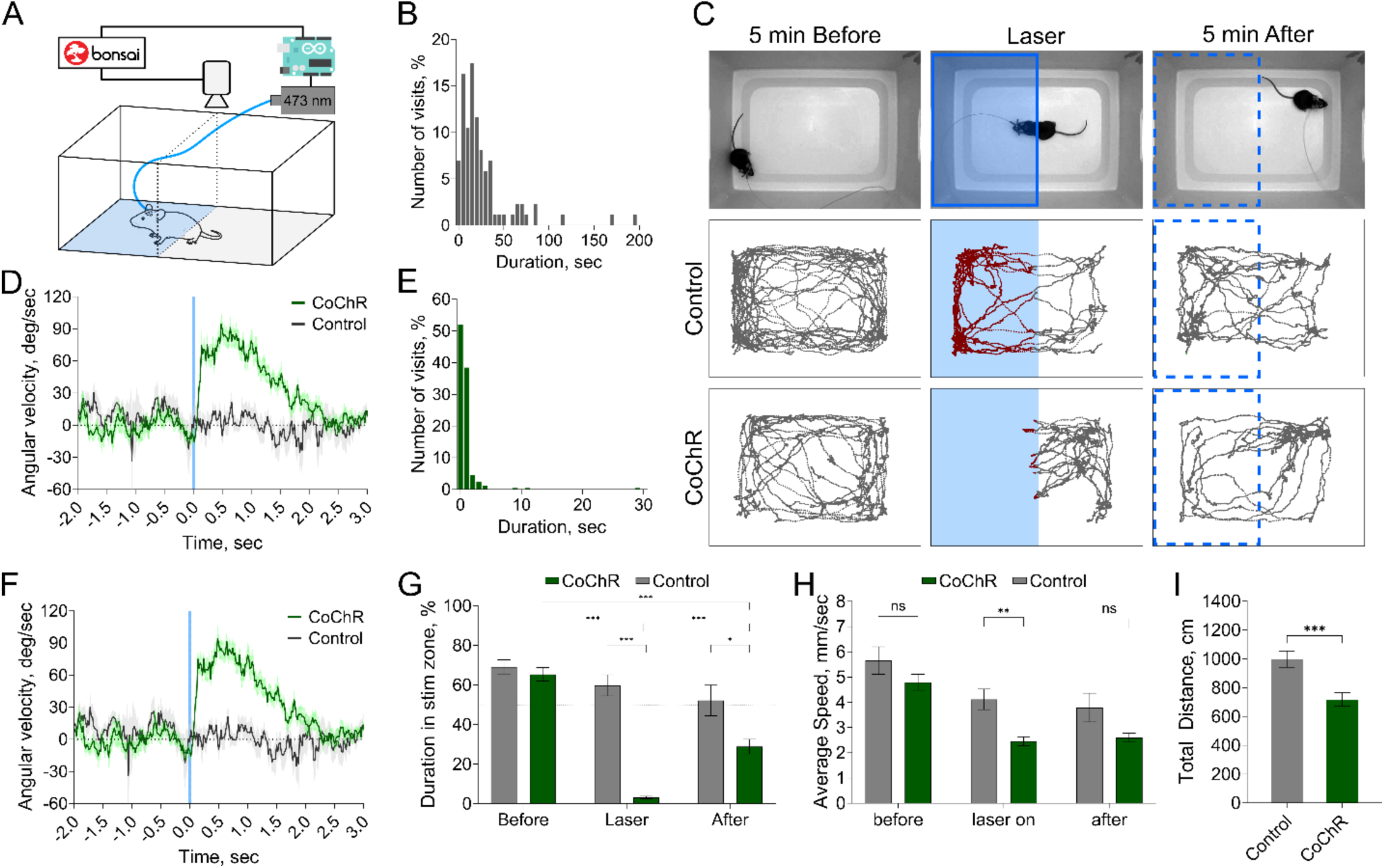
Optogenetic activation of Vglut2^+^-Pf neurons is aversive. A) Scheme of the setup for real-time place avoidance. B) Distribution of the duration of the visits in Control group during the ‘Laser’ epoch. C) Protocol of the real-time place avoidance. D) Instant body speed aligned to the onset of the laser during RTPA. Blue line – laser onset. E) Distribution of the duration of the visits in CoChR group during the ‘Laser’ epoch. F) Angular velocity of body to head vector aligned to the onset of the laser during RTPA. G) Time spent in the Laser—assigned half of the arena as a percent from the 5-minute epoch period. H) Average body speed in ‘Safe’ compartment during laser presentation and during ‘After’ period. I) Total distance during the ‘Laser’ epoch. All data represented as mean ± SEM, * - p< 0.0332, ** - p < 0.0021, **** - p < 0.0001, ns – p > 0.05.

Movement kinematics in the “Laser” half mirrored those in the open-field experiment, with mice decreasing body speed and performing small ipsilateral turns immediately after laser onset (Figure 2D, F). The effects were less pronounced due to shorter stimulation durations (typically <1 second; Figure 2E) compared to the 2-second stimuli used in the open field. Remarkably, avoidance developed after very brief Pf stimulation periods (Figure 2B, E). While control mice spent extended periods in the “Laser” compartment (average visit length 33.2 ± 25.3 seconds), CoChR mice entered and received stimulation for much shorter durations (average visit length 1.6 ± 2.3 seconds, range 0.28–9.4 seconds).

Together, these results show that optogenetic activation of Vglut2⁺ Pf neurons is sufficient to produce negative valence, driving rapid avoidance of the stimulation-paired compartment.

### Optogenetic Excitation of Vglut2^+^-Pf Neurons Induces Changes in Mouse Facial Expression

Optogenetic activation of Vglut2^+^-Pf neurons in the same animals produced robust turning effects and reliable avoidance in RTPA assay. These distinct behavioral outputs led us to a hypothesis that activation of Vglut2^+^-Pf neurons engages coordinated physiological response that may be expressed differently depending on the task or context. In addition, we also measured orofacial movements and pupil dynamics in head-fixed mice, recording their faces at 100 fps during stimulation.

Stimulation produced rapid movements across various facial regions, not limited to the whiskers (Figure 3A). These changes involved whisker, whisker pad, and nose movements, pupil constriction, and, in some cases, eye movements and jaw movements. For further analysis, we focused on the most prominent effects, using DLC to track the position and speed of the whisker pad (marked at the base of a macrovibrissa), nose, and pupil diameter (Figure 3A, B). Immediately after laser onset, CoChR mice moved the whisker pad forward, continuing to sweep forward and backward at this protruded position throughout the stimulation (Figure 3C). The average speed of the whisker pad was significantly higher in CoChR mice than in control animals (one-way ANOVA, F_(3, 63)_ = 8.012, p = 0.0001; post hoc Tukey 2mW vs. Control p = 0.003; 4mW vs. Control p = 0.0008; 6mW vs. Control p <0.0001; Figure 3D), though it did not vary with increased laser power. No lateralization was observed, as both sides of the whisker pad moved simultaneously and at the same speed during unilateral stimulation (Figure 3E). The latency to the first whisker pad movement start was 29 ± 22 ms (Figure 5K), shorter than the turn initiation latency, suggesting that orofacial movements are recruited early relative to whole-body reorientation. The whisker pad movement pattern varied with stimulation frequency (Figure 3F). Active whisking persisted after the end of stimulation (Figure 3C), consistent with the body speed increase observed in the open-field experiment. Nose movements were similar, with increased speed during stimulation (Figure 3H).

**Figure 3.**
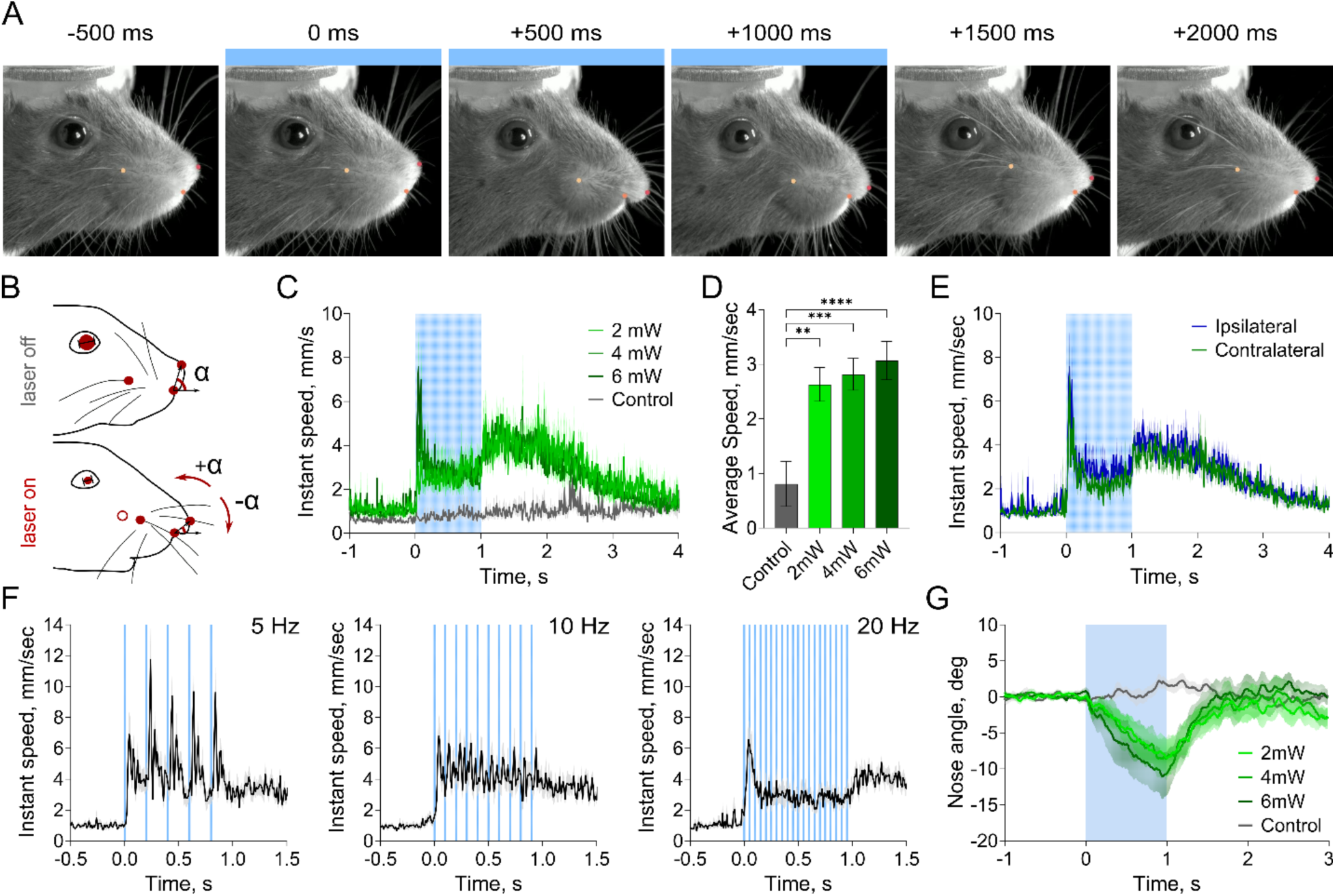
Excitation of Vglut2^+^ Pf neurons produces changes in facial expression. A) Example snapshots from CoChR animal representing changes in facial expression caused by optical excitation of Vglut2^+^-Pf neurons (10 ms pulses at 20 Hz, 6mW, blue line on snapshots). B) Schematic representation of main markers detected on the mouse face with DLC and calculated variables, such as whisker pad position and speed, nose position and pupil diameter. C) Instant speed of the whisker pad increases after laser onset in CoChR mice, but not in Control. D) Average speed of whisker movements didn’t change much with increase of the laser power. E) Instant speed of the whisker pad changes similarly on the ipsilateral and contralateral sides during unilateral stimulation. F) Whisker movements synchronized with the frequency of the laser pulses delivery. G) Nose angle decreases during laser presentation in CoChR mice but doesn’t change in Control mice. All data represented as mean ± SEM, * - p< 0.0332, ** - p < 0.0021, **** - p < 0.0001, ns – p > 0.05.

### Vglut2^+^-Pf Neurons Drive Both Somatic and Autonomic Responses

A notable finding from head-fixed recordings was the constriction of pupil diameter (Figure 4A, B), a response mediated by the parasympathetic nervous system via the smooth sphincter muscle of the iris. This pupil constriction we observed resembled the pupil light reflex (Hussain et al., 2009), characterized by a long latency, rapid constriction, and slower biphasic re-dilation (Figure 4A, B). We observed this pattern across various stimulation parameters with average latency 224 ± 69ms (measured in session with 6mW, 20Hz, Figure 5K). In contrast, control animals exposed to the same laser stimulation showed no pupil constriction (Figure 4C, D), verifying that the response resulted from Vglut2^+^-Pf neuron activation, not from the response to the laser light stimulation. We measured both ipsilateral and contralateral pupils (Figure 4C, D) and found that the amplitude of constriction was greater on the contralateral side in CoChR mice, but not in Control (two-way ANOVA, for Virus effect F_(1, 29)_ = 80.71, p < 0.0001; for Side effect F_(1, 29)_ = 6.005, p = 0.02; Fisher’s LSD CoChR Ipsilateral vs. Contralateral p = 0.0003; CoChR n = 20 stimulation sites, Control n = 11 stimulation sites). Further analysis was performed on the data from contralateral pupil. The amplitude of constriction was smaller at 2 mW compared to 4 and 6 mW (RM ANOVA, Power effect F_(1.917, 30.67)_ = 28.93, p < 0.0001; post-hoc Tukey, 2 mW vs. 4 mW p < 0.0001, 2 mW vs. 6 mW p < 0.0001; Figure 4E, F). Also the amplitude of constriction increased significantly with higher laser frequency, with 5 Hz producing a smaller effect than 10 or 20 Hz (RM ANOVA, Frequency effect F_(1.487, 28.25)_ = 33.36, p < 0.0001; post-hoc Tukey, 5 Hz vs. 10 Hz p < 0.0001, 5 Hz vs. 20 Hz p < 0.0001, 10 Hz vs. 20 Hz p = 0.03; Figure 4G, H). The effects of Vglut2^+^-Pf neuron activation on the ipsilateral pupil have similar patterns (Figure 4 I-L).

**Figure 4.**
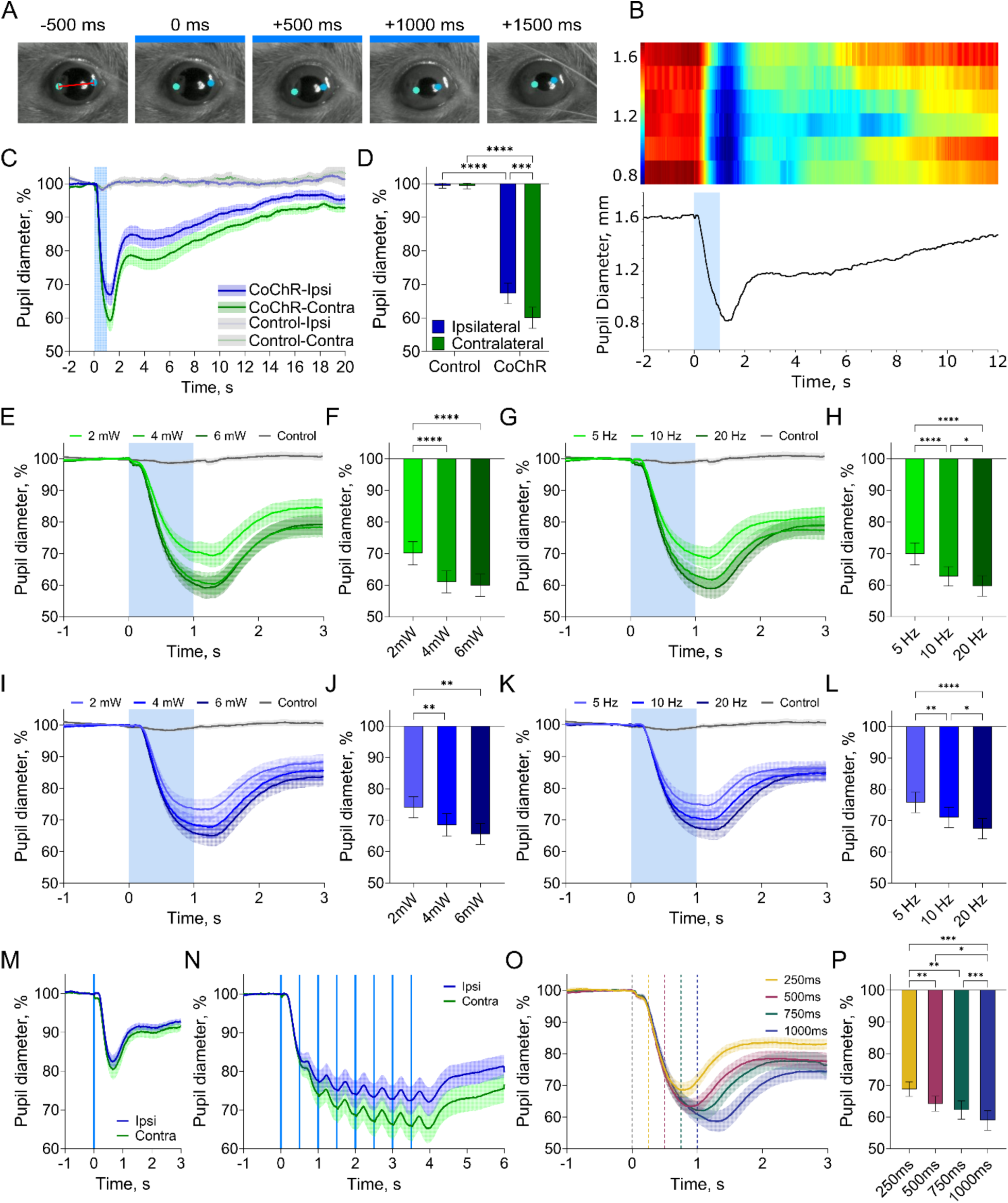
Excitation of Vlgut2^+^ Pf neurons causes pupil constriction. A) Example snapshots from CoChR animal before, during (snapshots with blue line) and after laser stimulation (10 ms pulses at 20 Hz, 6mW). Dots – deeplabcut markers for the pupil nasal and caudal border detection, red line – representation of pupil diameter. B) Representative pupil diameter changes in CoChR mouse during one session (6mW laser power, train of 10ms laser pulses at 20 Hz frequency). Up – heatmap represents pupil diameter change in 6 different trials, down – average change of the pupil diameter. C) Pupil diameter decreases during laser stimulation (6mW, 20Hz) period in CoChR, but no in control mice. Data normalized to the 1 second before stimulation. D) Average change of the pupil diameter as in C, calculated for the first 200ms after the laser end period. In CoChR mice the change in pupil diameter is higher in the contralateral side, then in ipsilateral. E) Stimulation at different laser power produces similar decrease of the contralateral pupil diameter. F) Average pupil diameter changes in 200ms after the laser end period. G) Stimulation at different laser frequencies produces similar decrease of the contralateral pupil diameter. H) Average pupil diameter changes in 200ms after the laser end period. I) Stimulation at different laser power produces similar decrease of the ipsilateral pupil diameter. J) Average pupil diameter changes in 200ms after the laser end period. K) Stimulation at different laser frequencies produces similar decrease of the ipsilateral pupil diameter. L) Average pupil diameter changes in 200ms after the laser end period. M) Single 10ms pulse stimulation at 6mW causes constriction pattern in both ipsilateral and contralateral pupils. N) 2Hz stimulation reveals the slow dynamic of the pupil response. O) Stimulation with increasing number of pulses (increasing duration) produces gradual decrease of the contralateral pupil diameter. P) Average pupil diameter changes in 200ms after the laser end period. All data represented as mean ± SEM, * - p< 0.0332, ** - p < 0.0021, **** - p < 0.0001, ns – p > 0.05.

In contrast to whisker movements time-locked to individual laser pulses (up to 10 Hz), the pupil response changes on a slower timescale. For further characterization of the temporal structure of this response, we recorded pupil dynamic in response of the stimulation consisted of different pulse numbers in a subset of CoChR mice (n=5). Even a single pulse (10ms long) elicited the similar constriction response started at 224 ± 26ms and reached the maximum constriction by 19.6 ± 6.9 % from baseline at 679 ± 109 ms (Figure 4M). Based on these latencies we next applied 2Hz stimulation protocol (8 pulses were delivered over 4 seconds), so laser pulses occurred close to the peak effect of the previous. Under this condition pupil constriction lasted for the duration of the stimulation and exhibited pulse-locked modulation (Figure 4N). Finally, we varied the number of pulses at fixed 20Hz frequency (250ms = 5, 500ms = 10, 750ms = 15 and 1000ms = 20 pulses) and showed that longer stimulation extended the constriction phase of the response resulting in larger response amplitude and longer latency to the maximum effect (Figure 4O). A repeated-measures ANOVA as a within-subject factor shown a significant main effect of the pulse number on maximum constriction amplitude (RM ANOVA, Pulse number effect F_(1.775, 14.20)_ = 28.35, p < 0.0001; post-hoc Tukey, 5 pulses vs. 10 pulses p = 0.0068, 10 pulses vs. 20 pulses p = 0.0166, 15 pulses vs. 20 pulses p = 0.0001, Figure 4P) and on time to maximum constriction (RM ANOVA, Pulse number effect F_(1.947, 15.58)_ = 71.94, p < 0.0001; post-hoc Tukey, 5 pulses vs. 10 pulses p = 0.0032, 10 pulses vs. 15 pulses p = 0.0173, 15 pulses vs. 20 pulses p = 0.0002). These results show that Pf activation can trigger pupil constriction with a single pulse, while the apparent pulse-locking depends on inter-pulse interval: if pulses are spaced by 500ms, individual pulses produce resolvable modulations, whereas with shorter gaps (e.g. 200ms at 5Hz) the pupil response becomes a smooth, low-pass–filtered constriction consistent with the slow dynamics of parasympathetic/smooth-muscle effector pathways

Altogether results from head-fixed experiment show that Vglut2^+^-Pf neuron stimulation evokes both the rapid whisker movements and slower pupil constriction with distinct temporal profiles indicating concurrent engagement of motor and autonomic output pathways.

### Pupil Constriction caused by Vglut2^+^-Pf Neuron Excitation Is Accompanied by Heart Rate Decrease

Given the pupil constriction observed in head-fixed mice, which is typically associated with parasympathetic activation, we investigated whether Pf stimulation could also produce additional parasympathetic effects. We recorded electrocardiogram (ECG) in freely moving mice (CoChR n = 6, Control n = 4) during optogenetic activation of Vglut2^+^-Pf neurons (Figure 5A). Electrodes were implanted on the rib cage above and below the heart, connected to a custom-made acquisition system and Blackrock. Offline analysis involved detecting R peaks and calculating instantaneous heart rate (HR) based on R-R intervals (Figure 5B). We found that HR decreased noticeably 1.5–2 seconds after stimulation onset (Figure 5C). To characterize this effect, we compared heart rate changes across different laser power and frequency conditions. Stimulation at 4 and 6 mW and 20Hz resulted in clear reduction in heart rate relative to control trials, while 2mW stimulation didn’t differ from control trials (Figure 5D, E). The strength of the heart rate change was higher at 6mW then at 4 mW. A repeated-measures ANOVA shown a significant effect of laser power on heart rate change (RM ANOVA, F_(1.734, 19.07)_ = 45.58, p <0.0001, post-hoc Tukey, 2 mW vs. 4 mW p < 0.0001, 2mW vs. 6 mW p < 0.0001, 4mW vs. 6mW p = 0.0429). Among the tested frequencies (5, 10 and 20Hz) only stimulation at 20 Hz produced a prominent decrease in heart rate relative to the control (Figure F, G). A repeated-measures ANOVA shown a significant effect of laser power on heart rate change (RM ANOVA, F_(1.471, 14.71)_ = 21.41, p =0.0001, post-hoc Tukey, 5Hz vs. 10Hz p = 0.0534, 5Hz vs. 20Hz p = 0.0011, 10Hz vs. 20Hz p = 0.003). To determine whether the amplitude of the heart rate decrease depended on the stimulation duration we varied the number of laser pulses with fixed power and frequency (CoChR n= 4, 6mW, 20Hz, Figure H-J). A repeated-measures ANOVA as a within-subject factor shown a significant main effect of the pulse number on maximum heart rate decrease amplitude (RM ANOVA, Pulse number effect F_(1.437, 10.06)_ = 12.03, p = 0.0036; post-hoc Tukey, 5 pulses vs. 10 pulses p = 0.0692, 10 pulses vs. 15 pulses p = 0.0291, 15 pulses vs. 20 pulses p = 0.9963, Figure 5I) and on time to minimum (RM ANOVA, Pulse number effect F_(1.750, 12.25)_ = 26.19, p < 0.0001; post-hoc Tukey, 5 pulses vs. 10 pulses p = 0.2732, 10 pulses vs. 15 pulses p = 0.0004, 15 pulses vs. 20 pulses p = 0.0289, Figure 5J). Interesting that post hoc comparison shown that stimulation durations of 750ms (15 pulses) and 1000ms (20 pulses) did not significantly differ from each other, suggesting the ceiling effect in the heart rate reduction.

Together, these findings demonstrate that optogenetic activation of Vglut2^+^-Pf neurons is sufficient to induce bradycardia and pupil constriction, indicating the engagement of parasympathetic output pathways.

### Motor and autonomic outputs in relation to Pf stimulation site

Recent studies indicate that Pf-neurons located in medial or lateral parts of the nucleus differ in their afferent and efferent connectivity and electrophysiological properties (Mandelbaum et al., 2019, Gonzalo-Martin et al., 2024). To evaluate the possibility that different behavioral effects were caused by stimulation of different anatomical parts of the Pf we examined the correlation between fiber position and the magnitude of the effects observed during stimulation (6mW power and 20Hz frequency). No significant correlation was found between location of the fiber tip along the mediolateral axis and average angular velocity during turn (R^2^ = 0.0085, p = 0.7, Pearson correlation), pupil diameter change (R^2^ = 0.0099, p = 0.677, Pearson correlation), or amplitude of the heart rate decrease (R^2^ = 0.0512, p = 0.4794, Pearson correlation, Figure 6 E-G). In addition, we divided CoChR stimulation sites into three groups based on the fiber position on the anterior-posterior axis: anterior (approximately AP -2.18,), middle (approximately AP -2.3) and posterior (approximately AP -2.46) and compared effects between these groups (Figure 6 A-D). Angular velocity differed significantly across Pf subregions (F_(2,17)_=4.07, p=0.036) with posterior sites producing larger turning amplitudes than anterior sites (post-hoc Tukey, p= 0.032). In contrast, neither pupil constriction nor heart rate decrease differed significantly between different stimulation sites (one-way ANOVA, pupil: F_(2,17)_=0.5538, p=0.58; heart rate: F_(2,9)_=0.2922, p=0.75).

**Figure 5.**
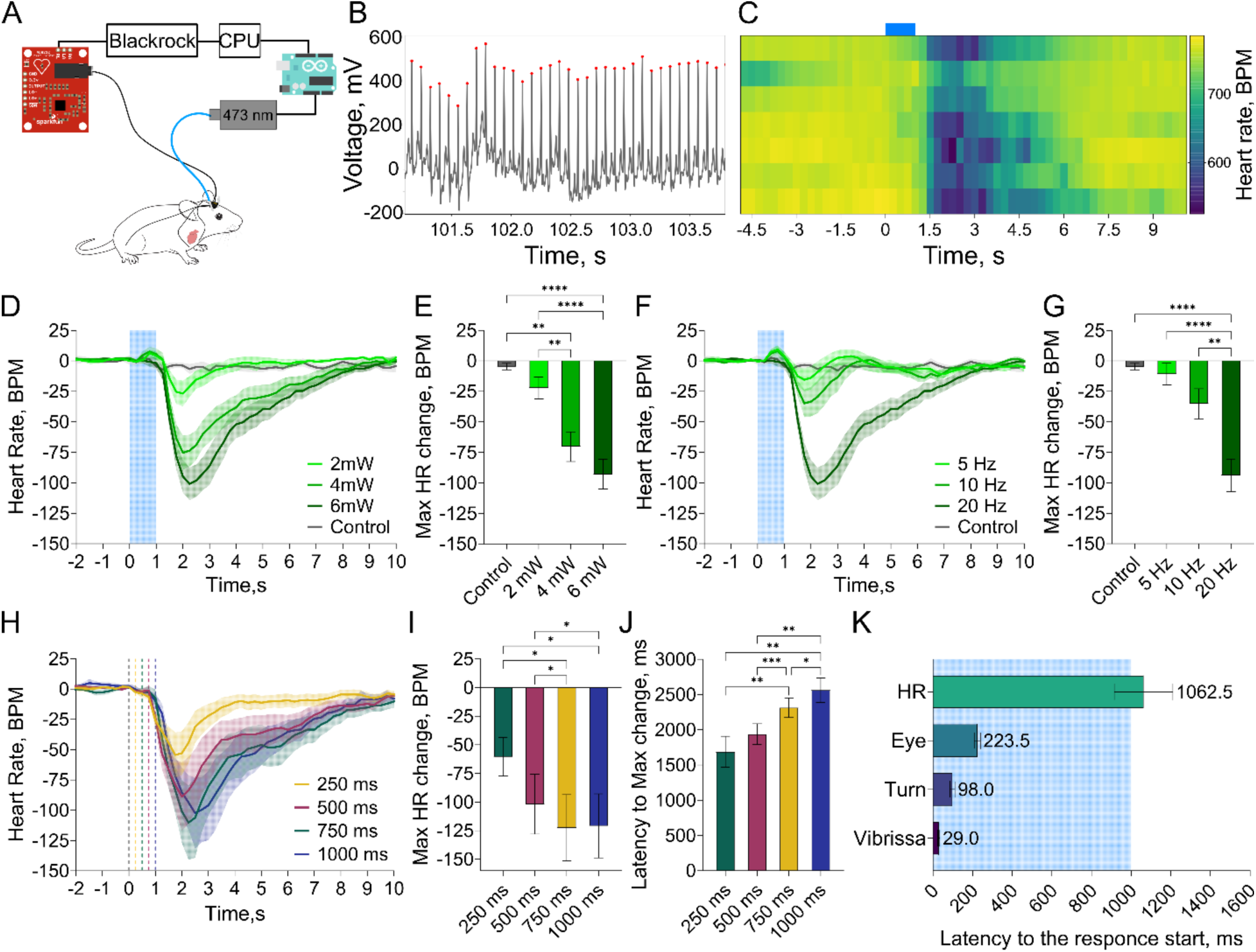
Optogenetic excitation of Vglut2^+^ Pf neurons reduces heart rate. A) Scheme of the ECG recording experiment. B) Representative example of raw ECG signal. Red dots – detected peaks, putative R peaks of ECG. C) Representative example of heart change rate following laser stimulation (blue rectangle, 10ms pulses at 20 Hz for 1 second, 6 mW). D) Instant heart rate decreases after the laser presentation. E) Maximal heart rate change occurs after laser presentation at 6 mW poser. F) Heart rate change depends on the frequency of the stimulation. G) Maximal heart rate change followed stimulation with 20 Hz frequency. H) Stimulation with increasing number of pulses produces gradual decrease of the heart rate. I, J) Latency and amplitude of heart rate change dependent on the duration of the laser stimulus. K) Skelemotor and autonomic responses appears at different latencies after the Vglut2^+^-Pf neuron stimulation. All data represented as mean ± SEM, * - p< 0.0332, ** - p < 0.0021, **** - p < 0.0001, ns – p > 0.05.

**Figure 6.**
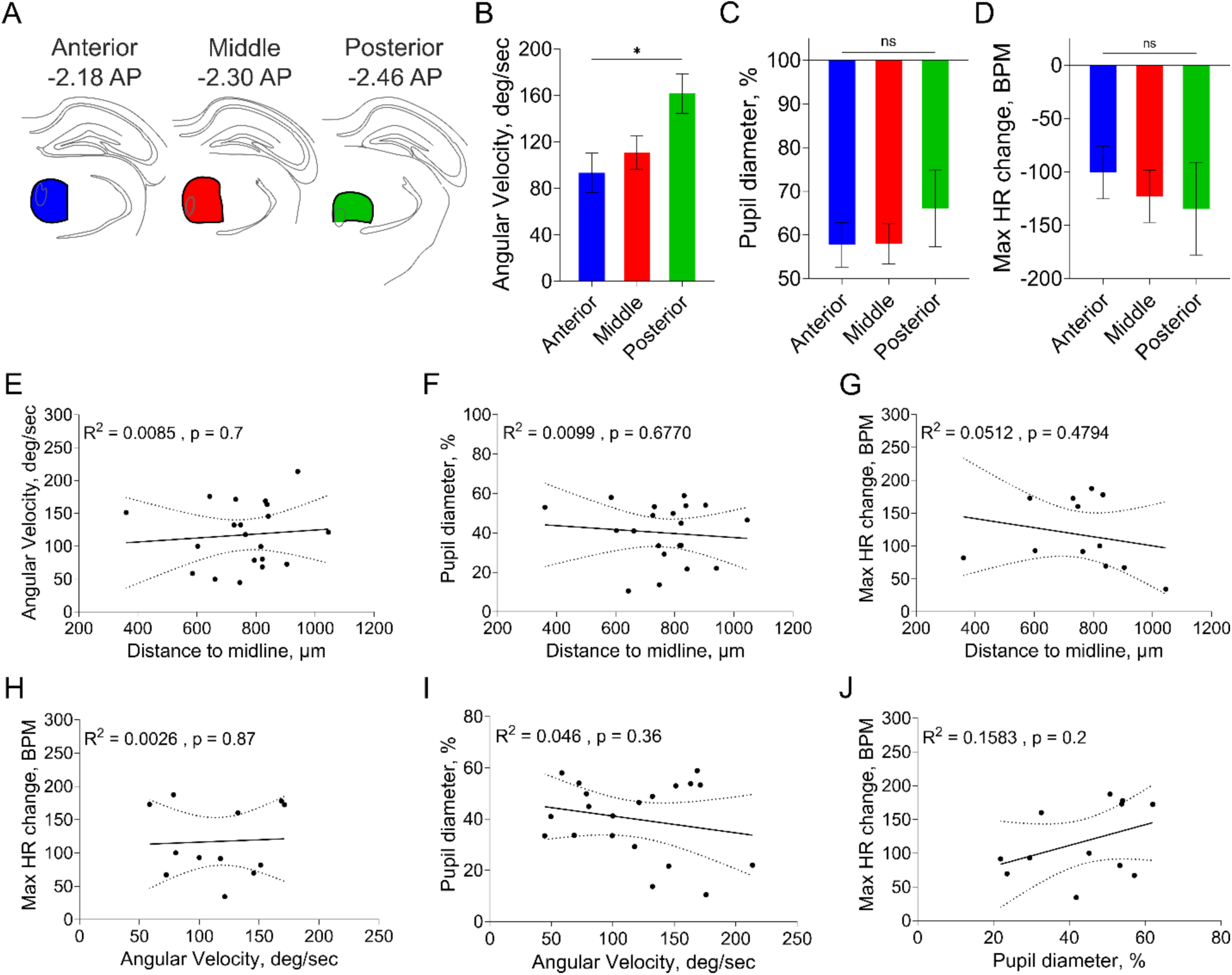
Anatomical and behavioral covariation of stimulation-evoked outputs within the parafascicular nucleus. A) Scheme of division Pf into three regions on the anterior-posterior axis. B-D) Anterior-posterior differences in average angular velocity (B), magnitude of pupil constriction(C) and maximal heart rate change amplitude (D). Bars represent mean mean ± SEM. E-G) Scatterplots showing the relationship between the mediolateral coordinate of the stimulation sites and the magnitude of average angular velocity (E), pupil diameter change (F) and heart rate change (G). H-J) Pairwise comparison of response amplitudes across all stimulation sites reveals no significant correlation between angular velocity and pupil diameter (H), angular velocity and heart rate(I) or pupil and heart rate (J) responses. Dots represent individual stimulation sites. * - p< 0.05

Because we recorded different behavioral and physiological outputs in the same animals, we next asked whether the magnitudes of these responses (angular velocity, pupil constriction and heart rate reduction) covaried across stimulation sites. We did not detect significant correlations between the different responses (p > 0.1 for all, Pearson correlation, Figure 6H-J).

Together, these analyses indicate that while posterior Pf stimulation sites bias the magnitude of turning behavior, autonomic outputs such as pupil constriction and heart rate reduction show no strong dependency on fiber position and do not covary with motor output magnitude.

## Discussion

In agreement with prior work, optogenetic excitation of Vglut2^+^ Pf projection neurons reliably induced fast ipsilateral turning (Watson et al., 2021; Fallon et al., 2023). Detailed kinematic analysis revealed a dissociation between head and body motion, with rapid head rotation leading changes in the center of mass. Pf has been proposed to relay vestibular information. Although we did not measure vestibular signals, the rapid head-first reorientation is consistent with Pf influencing orienting circuits that integrate vestibular and postural information(Smith et al., 2025). Together, these findings indicate that Pf activation coordinates skeletomotor and parasympathetic responses.

Beyond turning, Pf activation also produces orofacial movements. In head-fixed mice, optogenetic stimulation triggered rapid whisker pad and nose movements with short latency. These responses were bilateral even with unilateral stimulation, in contrast to the lateralized turning behavior, which is distinct from the effects of unilateral stimulation on turning. The bilateral nature of orofacial responses may reflect Pf’s bilateral projections or convergence onto midline brainstem motor circuits.

A striking and unexpected finding was significant pupil constriction following Pf activation. Pupil diameter is often used as a readout of general arousal level (McCormick et al., 2020). In mice the most constricted pupil is usually found during deep sleep (Yüzgeç et al., 2018), while locomotion is associated with pupil dilation, which depend on noradrenergic and cholinergic signaling (Reimer et al., 2016). In contrast, Pf activation produced pupil constriction despite increased motor output. Pupil constriction is classically observed in the pupillary light reflex which adjusts the amount of the light entering the eye. Because the constriction is performed by activation of smooth sphincter muscles, the latency is quite long, e.g. 246.4 ms in rats (Grozdanic et al., 2002) and 300 ms in mice (Hussain et al., 2009). The pupil constriction we detected also has a long latency of 224 ± 69 ms, suggesting a similar effector mechanism as the pupillary light reflex. Constriction was greater contralaterally and scaled monotonically with laser power, frequency and stimulation duration, consistent with gradual recruitment of a parasympathetic effector pathway.

Pf stimulation also induced a delayed decrease in heart rate, another indication of parasympathetic activation. The long latency and parameter dependence of pupil constriction and bradycardia suggest recruitment of parasympathetic effector pathways rather than nonspecific arousal. Pf activation seems to produce a coordinated state shift distinct from canonical sympathetic fear responses. Previous work has shown that some Pf neurons respond to vagal input and project to the striatum (Ito and Craig, 2005; Ito and Craig, 2008), and that Pf is interconnected with the periaqueductal gray, a key node in autonomic and defensive control (Sakata et al., 1988).

Pf activation was aversive in an RTPA task, consistent with earlier reports implicating Pf in avoidance-related behaviors (Delacour, 1969; Quiroz-Padilla et al., 2007) and with recent work identifying Pf subpopulations involved in encoding emotional valence (Chen et al., 2025; Li et al., 2025). However, the physiological profile observed here differs from canonical fear states, which are typically associated with sympathetic activation and pupil dilation. The dissociation between negative valence and sympathetic activation suggests that Pf-induced aversion may reflect destabilization of internal postural and autonomic set points rather than classical threat processing. This interpretation aligns with Pf’s anatomical connections to brainstem and midline thalamic nuclei involved in state regulation.

The temporal structure of the stimulation-evoked responses suggests hierarchical recruitment of distinct effector systems (**Figure 5**). Orofacial movements emerged first (whisker pad latency ∼29 ms), followed by whole-body reorientation (∼100 ms), indicating rapid engagement of fast motor circuits, likely via descending projections influencing brainstem and basal ganglia–motor pathways. Autonomic responses followed on a slower timescale: pupil constriction began at ∼224 ms, consistent with recruitment of brainstem parasympathetic nuclei and their preganglionic projections, while bradycardia emerged even later (∼1.5–2 s), reflecting additional peripheral dynamics of cardiac vagal modulation. The progressively increasing latencies, from skeletal muscle activation to smooth muscle control of the pupil and cardiac modulation, are consistent with hierarchical effector recruitment rather than a single unified motor command. This temporal cascade supports the interpretation that Pf activation triggers a coordinated state transition engaging both rapid motor outputs and slower parasympathetic-associated autonomic pathways.

Previous studies have demonstrated anatomical and functional heterogeneity within the Pf, with medial–lateral and anterior–posterior subdivisions differing in connectivity and function. (Mandelbaum et al., 2019; Gonzalo-Martín et al., 2024). While posterior Pf stimulation produced faster turning, autonomic responses were consistent across stimulation sites and did not covary with turning magnitude. This pattern suggests that autonomic and skeletomotor outputs of Pf activation are at least partially dissociable and not organized along a clear topographic gradient.

A limitation of the present study is that optogenetic stimulation was not restricted to projection-defined Pf subpopulations; nor did we assess necessity using inhibitory manipulations. However, our goal was to characterize the behavioral and physiological consequences of activating Vglut2⁺ Pf neurons as a population across contexts. This approach allowed us to reveal coordinated motor, autonomic, and motivational responses and their temporal structure, providing a functional framework that can guide future projection-specific and loss-of-function studies.

Collectively, the present findings suggest that Pf contributes to rapid state transitions that coordinate orienting movements, autonomic tone, and motivational valence. Rather than encoding motor commands per se, Pf activation may bias the organism into a specific behavioral state characterized by ipsiversive orienting and parasympathetic engagement. These findings have potential implications for therapeutic strategies targeting Pf circuitry. Pf has been considered as a target for deep brain stimulation in Parkinsonian conditions (Kerkerian-Le Goff et al., 2009) and autonomic dysfunction is increasingly recognized as a significant component of Parkinson’s disease (Kincl et al., 2024). Activation of Pf-STN projections can also rescue Parkinsonian akinesia. Our results suggest that stimulation parameters and pathway specificity will be critical determinants of therapeutic outcome, as indiscriminate Pf activation may engage autonomic circuits alongside motor pathways.

## Acknowledgment

This study is supported by NIH R01NS121253 to HHY. We would like to thank Gary Lehew for help with ECG recordings, and Ofer Yizhar for providing the plasmids for soma-targeted CoChR.

